# Differential encoding of noxious heat and self-reported pain along corticospinal networks: a simultaneous spinal cord-brain fMRI study

**DOI:** 10.1101/2025.10.24.684476

**Authors:** Dario Pfyffer, Yiyu Wang, Merve Kaptan, Joel Fundaun, Troy C Dildine, Valeria Oliva, Teresa Indriolo, Stefan Skare, Tim Sprenger, Philip K Lee, Minh Truong, Kenneth A Weber, Gary H Glover, Christine S W Law, Sean Mackey

**Affiliations:** Stanford University School of Medicine, Department of Anesthesiology, Perioperative and Pain Medicine, Palo Alto, United States; Center for Behavioral Sciences and Mental Health, Italian National Institute of Health, Rome, Italy; Department of Clinical Neuroscience, Karolinska Institutet, Stockholm, Sweden; MR Applied Science Laboratory Europe, GE Healthcare, Stockholm, Sweden; Stanford University School of Medicine, Department of Radiology, Stanford, United States

**Keywords:** corticospinal imaging, simultaneous brain and spinal cord fMRI, nociceptive processing, noxious heat, intensity encoding, self-reported pain, functional connectivity, representational similarity analysis

## Abstract

Chronic pain poses a substantial public health burden. Elucidating how the healthy central nervous system (CNS) differentially encodes objective stimulus intensity and subjective experiences of pain perception may offer key insights into the central mechanisms contributing to chronic pain. Functional MRI (fMRI) combined with controlled noxious stimulation provides a powerful means to explore neural representations of nociception and pain perception. Here, we applied noxious heat at three intensities (46 °C, 47 °C, 48 °C, 8 trials each randomized) to the right forearm of 28 healthy women during simultaneous spinal cord–brain fMRI to investigate how distributed corticospinal activity and connectivity encode stimulus intensity and subjective pain. Activity increased with stimulus temperature across regions involved in pain processing—including somatosensory, motor, prefrontal, insular, and subcortical areas—as well as in the ipsilateral dorsal and ventral spinal cord. Spinal– brain functional connectivity was observed between the right dorsal horn and pain-related brain regions such as primary and secondary somatosensory cortex, insula, anterior cingulate cortex, thalamus, and periaqueductal gray, and was positively associated with individual pain ratings. Using representational similarity analysis (RSA), we found that multivoxel activation patterns in the brain and spinal cord, as well as corticospinal connectivity patterns, reliably tracked stimulus temperature, while only subsets of cortical regions (e.g., insula, sensorimotor, and frontal cortices) encoded subjective pain. Notably, spinal cord representations were primarily organized by stimulus temperature rather than perceived pain intensity. These findings demonstrate that simultaneous spinal cord–brain fMRI combined with multivariate modeling can identify sensory and perceptual components of nociceptive processing across the neuroaxis. Such approaches advance mechanistic understanding of pain and may inform the development of CNS-based biomarkers for chronic pain assessment and intervention.

## Introduction

Chronic pain is a globally prevalent and heterogenous condition often treated for its symptoms and not its underlying mechanisms^1,2^. A fundamental step towards developing effective pain treatments is a comprehensive understanding of the central nervous system (CNS) mechanisms underlying pain perception. Functional magnetic resonance imaging (fMRI) combined with experimental pain stimulation allows for non-invasive measurement of neural activation along key nociceptive pathways to examine the underlying neural representations of pain experiences^3,4^.

Noxious heat stimulation is a commonly used pain paradigm in fMRI research that reliably elicits neural activation in established pain-processing networks in the brain^5–8^. The evoked neural activity typically includes bilateral activation corresponding to distinct stimulus intensities^9–11^. More recently, spinal cord fMRI has provided complementary knowledge of CNS correlates of nociceptive processing^12^, adding a critical piece along the neuroaxis to fully understand mechanisms of pain perception. Using noxious heat stimulation applied to the upper limb, previous studies have mapped the induced cervical spinal cord activity predominantly in the ipsilateral dorsal quadrant at the spinal level related to the stimulated dermatome, where second-order sensory neurons reside within the gray matter (GM)^13–21^. Notably, the activity increased in magnitude and extent when comparing noxious to innocuous stimulation^20^. However, there is limited knowledge of how these regions differentially respond to stimulations of various noxious intensities, especially along corticospinal pathways.

Recent methodological innovations in fMRI enable simultaneous acquisition of brain and cervical spinal cord images^22,23^, allowing a better understanding of their intricate interplay during nociceptive processing. Combined corticospinal activity has been investigated using left-arm noxious heat stimulation at two different intensities^24^. Cervical spinal responses increased in magnitude and spatial extent at higher vs lower intensities, paralleled by enhanced activation in pain-processing regions of the brain. The authors have shown a correlation between spinal cord-brainstem coupling and perceived pain intensity. However, previous studies mostly applied univariate analysis techniques which only focus on the fMRI signal amplitude in response to different levels of pain^25^. Using multivariate pattern analysis, brain fMRI studies have demonstrated that pain is represented in distributed patterns of activity, spanning the primary somatosensory cortex (S1), prefrontal and limbic regions, insula, and thalamus, some of which more closely track objective stimulus intensity (i.e., nociception), while others more closely track subjective experiences of pain perception^26–28^. Yet, comparable multivariate approaches have not integrated the spinal cord and spinal cord-brain functional connectivity (FC) to investigate how distributed corticospinal activation and FC patterns encode stimulus intensity and self-reported pain.

Building on that, in this study we systematically examine corticospinal activity and connectivity across three noxious heat stimuli using simultaneous spinal cord-brain fMRI. We expand current brain-only multivariate models to the spinal cord to comprehensively characterize encoding of nociception and pain perception across the neuroaxis. Specifically, we utilize representational similarity analysis (RSA)^29^ to identify corticospinal networks representing various intensities of noxious heat and interindividual differences in self-reported pain. These advances in understanding of the multifaceted nature of pain neurophysiology and can guide the development of future pain biomarkers.

## Methods

### Participants

This study includes 28 healthy females (age: 25.6±5.6 years). The participants completed the fMRI scan at the Richard M. Lucas Center for Imaging in Stanford, California, USA. The inclusion criteria were: between the ages of 18 and 65 years, female sex, no current pain, no history of chronic pain, no neurological or psychiatric condition, no history of opioid or psychoactive substance use, and MRI safe. Before assessments, all participants provided informed written consent and were instructed about the scanning procedure and the task. The study was approved by the Stanford Institutional Review Board. This study is part of a larger study in fibromyalgia which affects mostly women, hence the inclusion of only females.

### Experimental design

The thermal intensity encoding paradigm used during the fMRI scan included noxious heat stimuli of fixed temperatures at 46 °C, 47 °C, and 48 °C. These were applied to the C6 dermatome at the right volar forearm using a 3×3 cm^2^ surface Advanced Thermal Stimulator (ATS) thermode connected to the PATHWAY (Medoc Ltd, Israel). The 46 °C, 47 °C, and 48 °C conditions consisted of eight blocks each which were pseudo-randomized. For each block, a ramp and hold sequence was used with a baseline temperature of 32 °C and a stimulation period of 12 s, including an 8 °C/s ramp up and 8 °C/s ramp down phase. This was preceded by jittered anticipation time of either 3.5, 4.5, or 5.5 s (pseudo-randomized) and followed by a pain rating period of 3.5 s. Participants were instructed to rate their subjective pain on a Visual Analog Scale (VAS) ranging from 0 to 10, 0 meaning no pain and 10 reflecting the worst pain imaginable (with mental anchors 3-weak pain, 5-moderate pain, and 7-strong pain). Pain was rated using a manually operated controller with buttons to move the cursor along the VAS projected onto a screen. A 9-second rest block followed each heat stimulation block, during which participants fixated on a plus sign (Fig. 1a). Visual cues were provided with Eprime (Version 2.0, Psychology Software Tools, Pittsburgh, PA). Pain ratings were extracted from the 0-10 VAS recordings in increments of 0.125.

**Figure 1.**
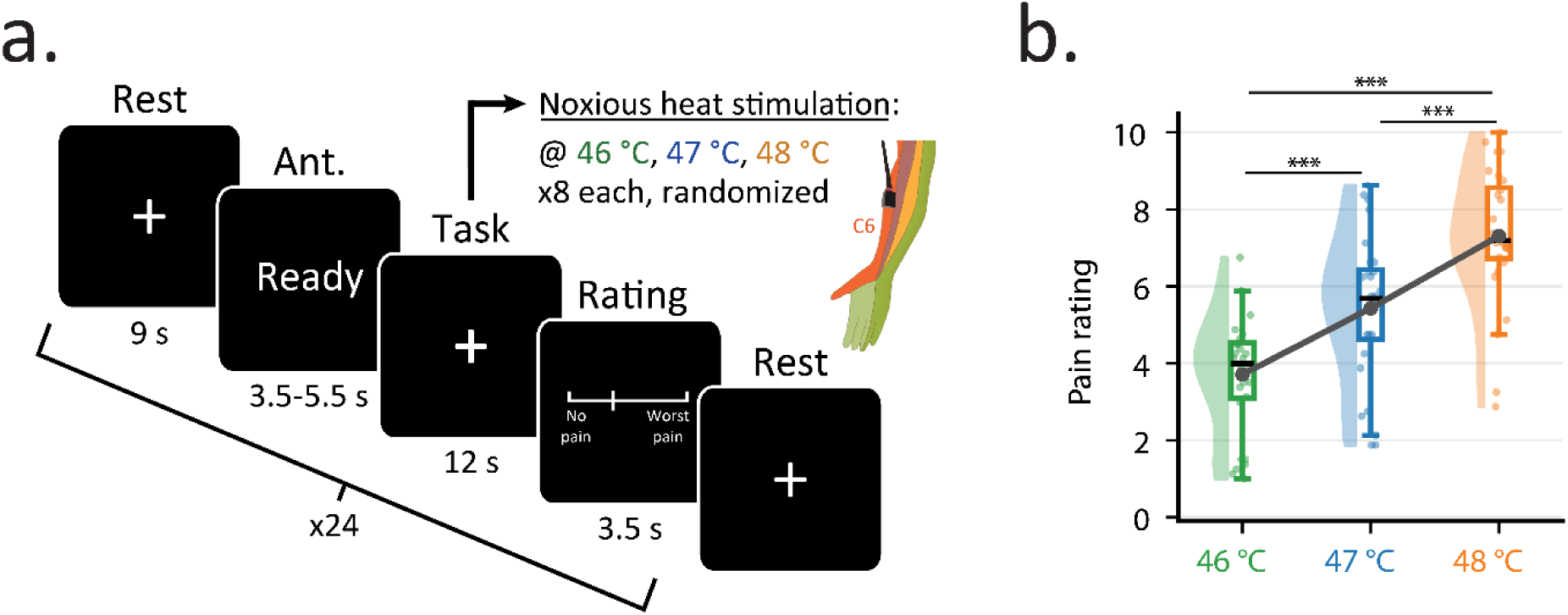
Experimental design. **a.** Each trial began with an anticipation (Ant.) period of 3.5, 4.5, or 5.5 seconds, indicated by the word “Ready” on the screen. Heat was then applied to the right volar forearm for a total of 12 seconds, followed by a 3.5-second pain rating period using a 0–10 visual analog scale (VAS: from no pain to worst pain imaginable) and a button box providing real-time visual feedback. A 9-second rest period concluded each trial. The sequence was repeated 24 times (8 trials per temperature). **b.** Pain ratings for each temperature are depicted via box plots. For the box plots, the median is denoted by the central mark, and the bottom and top edges of the boxes represent the 25th and 75th percentiles, respectively, with the whiskers encompassing ∼99% of the data. The circles represent individual participants, and half-violin plots show the distribution across participants. Gray circles (connected via lines) denote the mean value. *** p < 0.001.

### MRI data acquisition

fMRI data were collected using a 3T General Electric Discovery 750 scanner equipped with a 16-channel neurovascular array head/neck coil. Functional runs consisted of 290 (first 8 dummy volumes were removed during preprocessing) gradient-echo EPI volumes (acquisition time: 12:05 min) that were acquired with two field-of-views to optimize the acquisition for brain and spinal cord. The sequence was developed on the KS Foundation platform (https://ksfoundationepic.org/), enabling the use of a spectral-spatial RF pulse for brain excitation and a 2D echo-planar selective RF pulse for reduced-FOV excitation in the spinal cord. The brain FOV consisted of 31 axial slices covering the whole brain and the spinal cord FOV consisted of 12 slices covering vertebrae C4-C7 with the following parameters: EPI readout sequence with GRAPPA acceleration factor R = 2, FOV for brain/spinal cord = 22/8 cm, flip-angle = 80°, TE = 30 ms, TR = 2.5 s, matrix size = 64×64, readout BW = ±250 kHz, in-plane resolution: 1.25×1.25 mm^2^ and slice thickness = 5 mm without gaps. Dynamic per slice shimming^23^ determined the optimal shim setting for brain and spinal cord slices automatically.

Additionally, spinal cord high-resolution T2-weighted Sagittal 3-D CUBE (TR/TE = 2502/100 ms, flip = 90°; matrix = 512 × 512 × 72, posterior–anterior × inferior–superior × left–right; voxel = 0.47 × 0.47 × 0.70 mm³) and high-resolution T1-weighted sagittal 3D BRAVO (TR/TE/TI = 7.6/2.9/400 ms, flip = 11°; matrix = 186 × 256 × 256, left–right × posterior–anterior × inferior–superior; voxel = 1.0 × 0.86 × 0.86 mm³) images were obtained for registration and normalization purposes. Cardiac and respiratory traces were recorded during functional scans using a finger pulse oximeter and pneumatic below, respectively.

### Brain and spinal cord data preprocessing

All acquired images were converted to NIfTI using dcm2niix (v1.0.20181125). The processing of functional and structural images was done using in-house bash and MATLAB scripts using tools from Spinal Cord Toolbox (SCT, versions 5.6.0 and 6.3.0 [for spinal level templates])^30^, and the Oxford Center for fMRI of the Software Library (FSL, version 5.0).

#### Motion correction

For brain data, 3D rigid-body alignment (with FSL ‘*flirt’*) was performed using rigid-body realignment^31^ with spline interpolation.

For the spinal cord, a three-step motion-correction was performed using FSL’s ‘*flirt’* tool with spline interpolation. First, all volumes were averaged to create a mean image.

Then, a binary mask limited to the region around the spinal canal was manually generated based on this mean image. This mask was used to ensure that regions outside of the cord did not adversely impact the motion correction procedure. In the first step of motion correction, each volume was aligned to the middle volume of the functional run using 3D rigid-body realignment. In the second step, the process in step 1 was repeated, but the volumes were aligned to the average volume from step 1. Finally, 2D slice-wise realignment - using the mean image from step 2 of motion correction as the reference - was employed for each axial slice to correct for the non-rigid motion of the cervical spine and physiological motion from swallowing and respiration.

For both brain and spinal cord, the temporal signal-to-noise ratio (tSNR) was calculated by dividing the mean of the time-series by their standard deviation after motion correction (Supplementary Fig. 1).

#### Time-series denoising

For both the brain and the spinal cord, a RETROICOR approach^32^ was used to address physiological noise by creating slice-specific regressors, with a total of four regressors to account for cardiac and respiratory processes (2 for cardiac and 2 for respiratory). Note that physiological noise correction was employed before slice-timing correction, as slice-timing correction was shown to alter the synchronization between the acquisition times of each slice and physiological data^33^.

Additionally, we regressed out i) motion parameters (6 for brain, 3 for spinal cord), ii) motion outliers, and iii) CSF signal by extracting the mean time-series using CSF masks (obtained with FSL’s ‘*fast*’ for brain, manually drawn for spinal cord) for each region. Motion outliers were determined by dVARS (the root mean square difference between successive volumes^34^) as a metric implemented in the ‘*fsl_motion_outliers’* function of FSL.

#### Normalization

Brain and spinal cord sub-volumes were registered to the MNI152 and PAM50 standard template through a two-step process. For the brain, each participant’s EPI image was registered to their T1-weighted structural scan via linear registration with 7 degrees-of-freedom. Next, the T1-weighted image of each participant was registered to the 2 mm MNI152 standard space T1-weighted image, using a 12 degrees-of-freedom linear fit followed by linear registration in FSL. The two resulting registration fields were then combined and applied to the functional data, bringing them into the template space.

For the spinal cord, the initial step involved normalizing anatomical T2-weighted images to the template space, completed in three consecutive steps (‘*sct_register_to_template*’): (i) straightening the spinal cord based on the binary manual cord segmentation, (ii) aligning vertebral levels C2-C7 using automatically labelled vertebrae (generated by ‘*sct_label_vertebrae*’, with manual corrections as needed), and (iii) registering the anatomical images to the template with non-rigid, segmentation-based transformations. In the second step, the mean preprocessed functional image was registered to the T2-weighted PAM50 template^35^ using non-rigid transformations (‘*sct_register_multimodal’*), with the initial alignment based on the warping field obtained from the first step.

#### Smoothing

Prior to data analysis, brain and spinal cord data were smoothed with a 5 mm isotropic and 2 x 2 x 5 mm^3^ anisotropic full-width half maximum Gaussian kernels with FSL’s *SUSAN*, and SCT’s *sct_smooth_spinalcord*, respectively.

### Behavioral data analysis

For each temperature, within-participant average pain ratings were calculated. To calculate the differences in pain ratings between each temperature, within-participant average pain ratings were subtracted from each other (e.g., average rating for 47 °C minus average rating for 46 °C).

### Brain and spinal cord data analysis

General linear model (GLM) approaches were used for brain and spinal cord sub-volumes separately. For both sub-volumes, the first-level design matrix included the conditions “Ready”, “46”, “47”, “48”, and “Rating”. Each experimental design regressor was modeled by convolving boxcar functions with the canonical hemodynamic response function (HRF). The first-level blood oxygenation level dependent (BOLD) contrasts of interest were defined as: i) 46, ii) 47, iii) 48, iv) linear (48, 47, 46 with weights 1, 0, −1). A temporal high-pass filter with a 200 s cutoff period and pre-whitening using FILM^36^ were applied to first-level analyses.

Second-level analyses were carried out using a mixed-effects model with the contrast images that were obtained during first-level analyses to generate group-level activation maps (corrected for multiple comparisons using Gaussian Random Field [GRF] theory, z > 1.6 or z > 3.1 for spinal cord and brain, respectively).

Prior to calculation of average z-scores within the activated/deactivated regions and the number of activated/deactivated voxels, the subject-level maps were thresholded at z>2.3 and z<-2.3 for the brain and z>1.645 and z<-1.645 for the spinal cord, respectively. The pairwise differences were assessed with t-tests (Bonferroni-corrected).

### Seed-based functional connectivity between the spinal cord and the brain

FC between the spinal cord and the brain was assessed by using a seed-based approach. As a seed region, the right dorsal quadrant of the spinal cord was chosen (see Fig. 3). To capture activation patterns that may extend beyond the GM due to partial volume effects, spatial smoothing, and registration mismatches to the template, a broader ROI encompassing the entire right dorsal quadrant was defined rather than restricting it to the right dorsal horn only. This approach acknowledges the spatial imprecision inherent in spinal cord fMRI and ensures that signal relevant to task-evoked activity is not unintentionally excluded due to methodological limitations. For each participant, the time-series from the right dorsal quadrant was extracted at C5-C7 spinal levels and correlated with each brain voxel’s time-series. Similar to Sprenger et al.^24^, this was performed within a GLM analysis to account for task-induced effects (i.e., eliminate activation-induced correlations)^37^ by additionally including the task-based regressors to the model to isolate intrinsic spinal cord-brain coupling. These first-level maps were then supplied to second-level analyses which were carried out using a mixed-effects model to generate group level connectivity maps (corrected for multiple comparisons using GRF theory, z > 1.6, and cluster-level threshold p < 0.05). The FC strength between the spinal time-series and brain was then correlated with each participant’s mean pain rating.

### Representational similarity analysis

RSA is a multivariate technique that allows mapping between neural activity and behavioral measures based on a shared structure in their similarity (or distance) matrices^29^. By specifying such representation structures, RSA can disentangle whether the neural representation structure in the brain and the spinal cord is related to temperature or pain encoding.

We computed the representational similarity matrix (RSM) of heat stimulation by calculating the pairwise distance between each trial temperature using Euclidean distance. This yielded a 24×24 symmetric matrix. Similarly, we computed the representational space of the pain ratings. The temperature and the pain-rating RSMs represented the two behavioral representation models that we would relate to the neural RSMs in the next steps.

To examine both the local and distributed neural representation, we applied RSA to both the neural activation throughout the neuroaxis parcellations and spinal cord-brain functional connectivity.

For the activation-based RSA, then neural RSMs estimated pairwise similarities of neural representations across the 24 trials by calculating the Euclidean distance between activation patterns. The mean trial activation was estimated using a trial-wise design GLM, which included a separate boxcar regressor for each trial. All other steps (e.g., nuisance, motion regressors, HRF) are identical to the conditional GLM. The local mean activation patterns were extracted using a combination of different parcellations. For the brain, we identified subcortical areas from the Pauli atlas^38^ and cortical ROIs from the Schaefer atlas^39^, resulting in a total of 112 brain ROIs (12 subcortical + 100 cortical areas). For the brainstem, we identified the periaqueductal gray (PAG) and the rostral ventromedial medulla (RVM) using the Brainstem Navigator toolkit (https://www.nitrc.org/projects/brainstemnavig/). For the spinal cord, we created masks of for the four quadrants (right dorsal, right ventral, left dorsal, left ventral) by dividing the PAM50 cord template in SCT.

To examine how temperature and pain representations were encoded in each region, these similarity measures were then regressed onto models reflecting temperature similarity and subjective pain similarity. For example, if trials with the same temperature condition also evoked highly correlated multivoxel neural activity patterns, the regression would indicate that temperature strongly predicted neural similarity. Likewise, if trials with comparable pain ratings showed more similar activity patterns, this would suggest that neural representations were organized by subjective pain. Since all RSMs are symmetric, the lower triangles of the temperature, pain, and neural RSMs were vectorized. The vectors were submitted to two regression models to examine whether the neural RSMs can be explained by the temperature or the pain rating RSMs for each subject. The group-level inference is conducted using a one-sample t-test on the individual-level regression coefficients from the regression models. Bonferroni correction was used on the t-tests to control for multiple comparisons.

The functional connectivity-based RSA (Fig. 2) follows the same procedure with only one modification. The neural RSMs were conducted using seed-to-seed FC during the heat stimulation period in each trial. Concretely, we extracted 5 time points corresponding to the heat stimulation period for each trial (12 seconds with a 2.5-second TR) for each of the brain ROIs, two brainstem ROIs, and the four spinal cord quadrants. We then calculated the trial-wise connectivity matrix between a spinal cord quadrant to the brain and brain stem ROIs.

**Figure 2.**
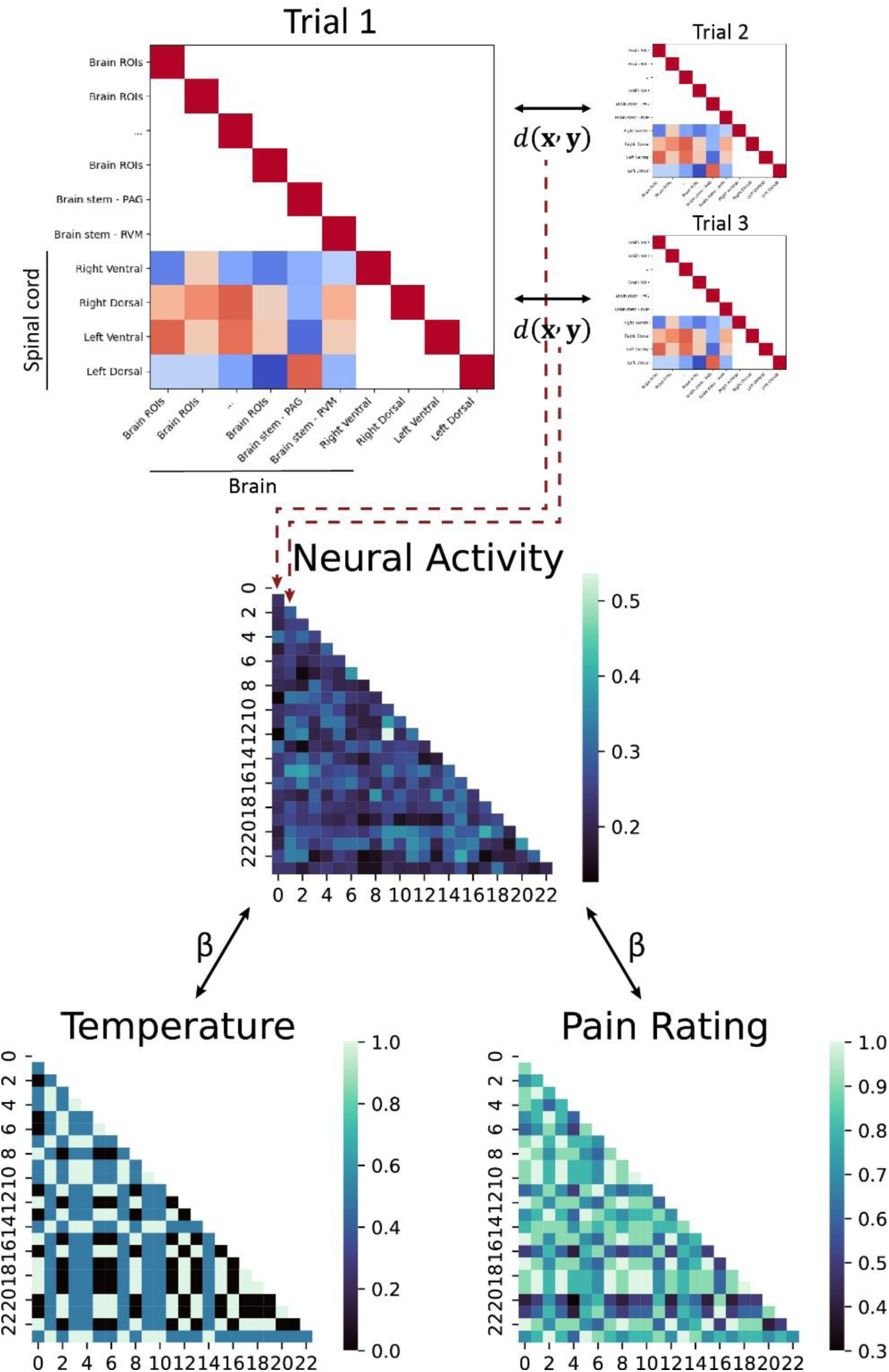
Schematic of representation similarity analysis for corticospinal functional connectivity. For each trial, seed-to-seed functional connectivity between spinal cord quadrants and brain regions during heat stimulation was computed, represented by the four colored rows in the lower triangle of the functional connectivity matrix for each trial (top panel). Pairwise Euclidean distances, d(x’ y), between connectivity matrices across all trial combinations were used to construct the neural activity representational similarity matrices (RSMs) (middle panel). Corresponding temperature and pain-rating RSMs were generated by calculating pairwise Euclidean distances between trials based on stimulus intensity and self-reported pain, respectively (bottom panel). The lower triangles of the temperature, pain, and neural RSMs were then vectorized and entered into separate linear regression models to determine whether neural representations reflected temperature or pain encoding.

Repeating this step for the four spinal quadrants resulted in four sets of connectivity-based neural RSMs. In the graphical representation (Fig. 2), the four rows in the lower triangle of a connectivity matrix of trial 1 capture the connectivity patterns from the four spinal cord quadrants to the brain and the brain stem. The pairwise distance between trials calculated used the entire row of values for the spinal cord-to-whole brain connectivity to capture the distributed functional connectivity patterns. In a follow up analysis to examine whether the behavioral patterns are represented by a specific pathway (i.e., ROI-to-ROI or network-to-network connectivity), the pairwise distance between trials were calculated used the corresponding values of the selected regions for the seed-to-seed connectivity. The distance metrics then are organized into a single neural RSM per participant. These neural RSMs were also submitted to regression models to examine how temperature and pain are encoded in connections along the neuroaxis.

## Results

### Self-reported pain increases with temperature

The mean pain rating for each temperature condition was 3.7±1.5, 5.4±1.9, and 7.3±1.8 for 46 °C, 47 °C, and 48 °C, respectively (Fig. 1b), demonstrating that the applied heat stimuli were perceived as painful and dissociable. Ratings increased by 1.7±1.0 from 46 °C to 47 °C (p < 0.001), by 1.9±0.8 from 47 °C to 48 °C (p < 0.001), and by 3.6±1.2 from 46 °C to 48 °C (p < 0.001).

### Neural activity increases with temperature in the brain and spinal cord

BOLD responses in the brain and spinal cord for each temperature against the baseline revealed a typical pattern of pain-processing regions reported with noxious heat stimulation (Fig. 3a, left panel). Consistent activation was observed across sensorimotor areas (S1, primary motor cortex [M1], premotor cortex [PMC], supplementary motor area [SMA]), dorsolateral prefrontal cortex [DLPFC], secondary somatosensory cortex [S2], insula, central opercular cortex, subcortical structures (pallidum, putamen, thalamus), cerebellum, and the brainstem (mixed effects, p < 0.001, cluster correction threshold p < 0.05). Deactivation was observed in default mode network (DMN) nodes such as PFC, cingulate gyrus, angular gyrus, middle frontal gyrus, and occipital cortex (mixed effects, p < 0.001, cluster correction threshold p < 0.05). Notably, the number of activated voxels increased significantly from 46 °C to 47 °C, 47 °C to 48 °C, and the linear contrast (Fig. 3a, right panel; p < 0.001). Similarly, the number of deactivated voxels also increased significantly from 47 °C to 48 °C (p < 0.05) and 46 °C to 48 °C (p < 0.01). The linear contrast showed widespread differences in activation and deactivation, with increased activity across sensorimotor cortices, ACC, insula, thalamus, and cerebellum, and decreased activity within ipsilateral S1, middle and inferior frontal gyri, lateral occipital cortex, paracingulate gyrus, and VLPFC.

**Figure 3.**
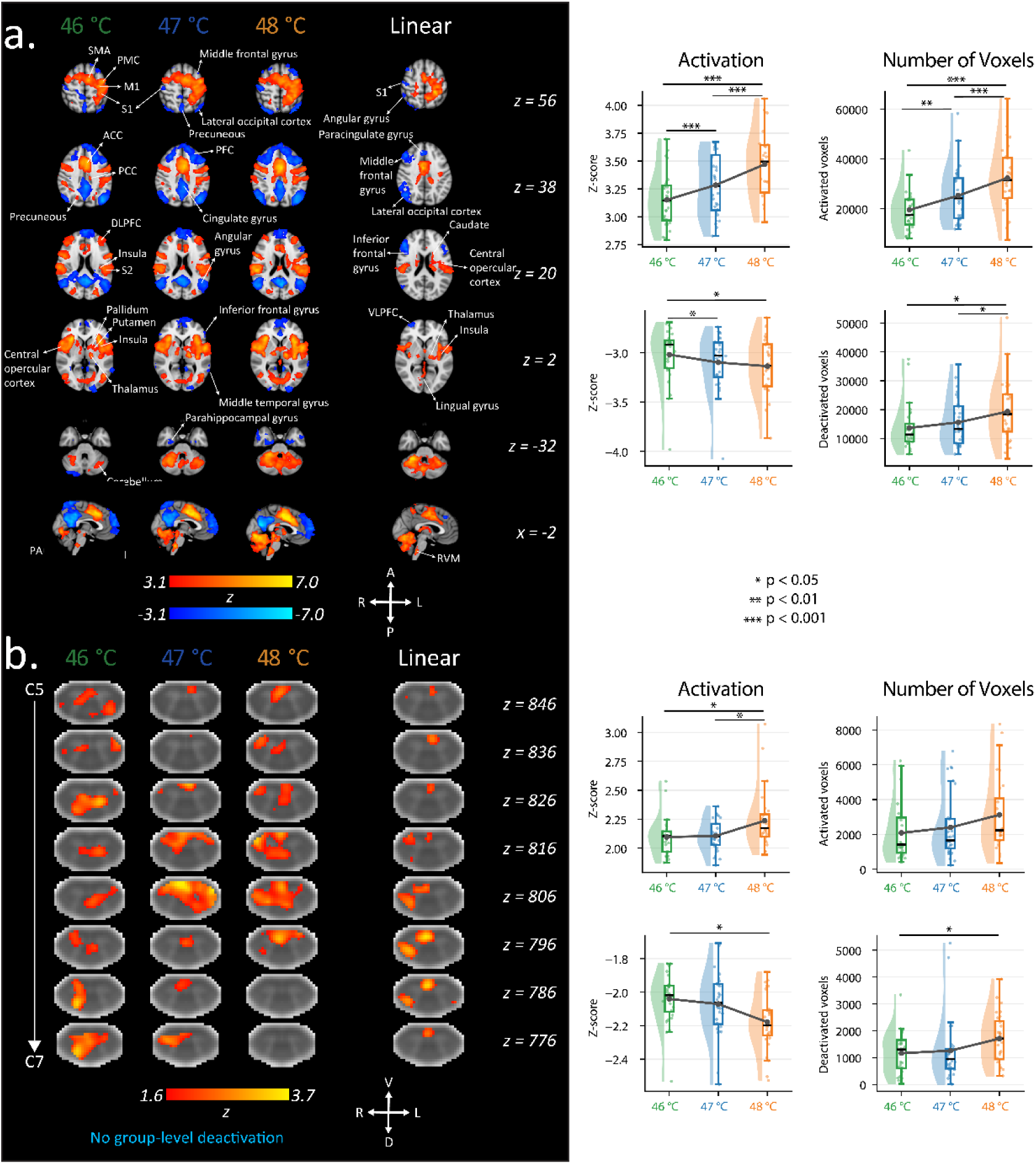
Self-reported pain ratings, brain and spinal cord activation at different heat stimulation intensities. **a**. Left panel. Group average activation maps (depicted on axial MNI152 slices, see the MNI coordinates on the right side) for the main effect of each temperature and the linear contrast (p < 0.05 FWE-corrected, mixed effects). Right panel. The average z-scores within the activated/deactivated regions, as well as the number of activated and deactivated voxels in the brain, are depicted via box plots. **b**. Left panel. Group average activation maps (depicted on axial PAM50 T2-star weighted axial slices, see the z-coordinates on the right side) for the main effect of each temperature (p < 0.05 FWE-corrected, mixed effects) and the linear contrast (p < 0.05 uncorrected, mixed effects). Right panel. The average z-scores within the activated/deactivated regions, the number of activated and deactivated voxels in the spinal cord are depicted via box plots. For all box plots, the median is denoted by the central mark and the bottom and top edges of the boxes represent the 25th and 75th percentiles, respectively, with the whiskers encompassing ∼99% of the data. The circles represent individual participants, and half-violin plots show the distribution across participants. Gray circles (connected via lines) denote the mean value. * p < 0.05, ** p < 0.01, *** p < 0.001. ACC, anterior cingulate cortex; DLPFC, dorsolateral prefrontal cortex; M1, primary motor cortex; PCC, posterior cingulate cortex; PFC, prefrontal cortex; PMC, premotor cortex; S1, primary somatosensory cortex; S2, secondary somatosensory cortex; SMA, supplementary motor area; VLPFC, ventrolateral prefrontal cortex.

Spinal cord activity maps showed distributed activation patterns covering dorsal and ventral horns across segmental levels C5-7 (Fig. 3b, left panel; mixed effects, p < 0.05, cluster correction threshold p < 0.05). At the group level, the number of active voxels increased across stimulation intensities; however, the difference was not statistically significant (p > 0.08). The linear contrast highlighted predominant activation of the ipsilateral dorsal horn and middle ventral spinal cord. In contrast, the number of deactivated spinal voxels was significantly higher at 48 °C compared to 46 °C (Fig. 3b, right panel; p < 0.05). No group level deactivation was observed with multiple comparison corrections.

### Functional connectivity between the spinal cord and the brain is modulated by subjective pain ratings

To examine FC between the spinal cord and the brain, the spinal cord time-series of the right dorsal quadrant was correlated with the time-series of all brain voxels (Fig. 4a, left panel). Before FC calculation, variance components associated with the experimental task regressors from the initial GLM were regressed out from the brain time-series to isolate intrinsic connectivity patterns that were not driven by mean signal changes^37^.

**Figure 4.**
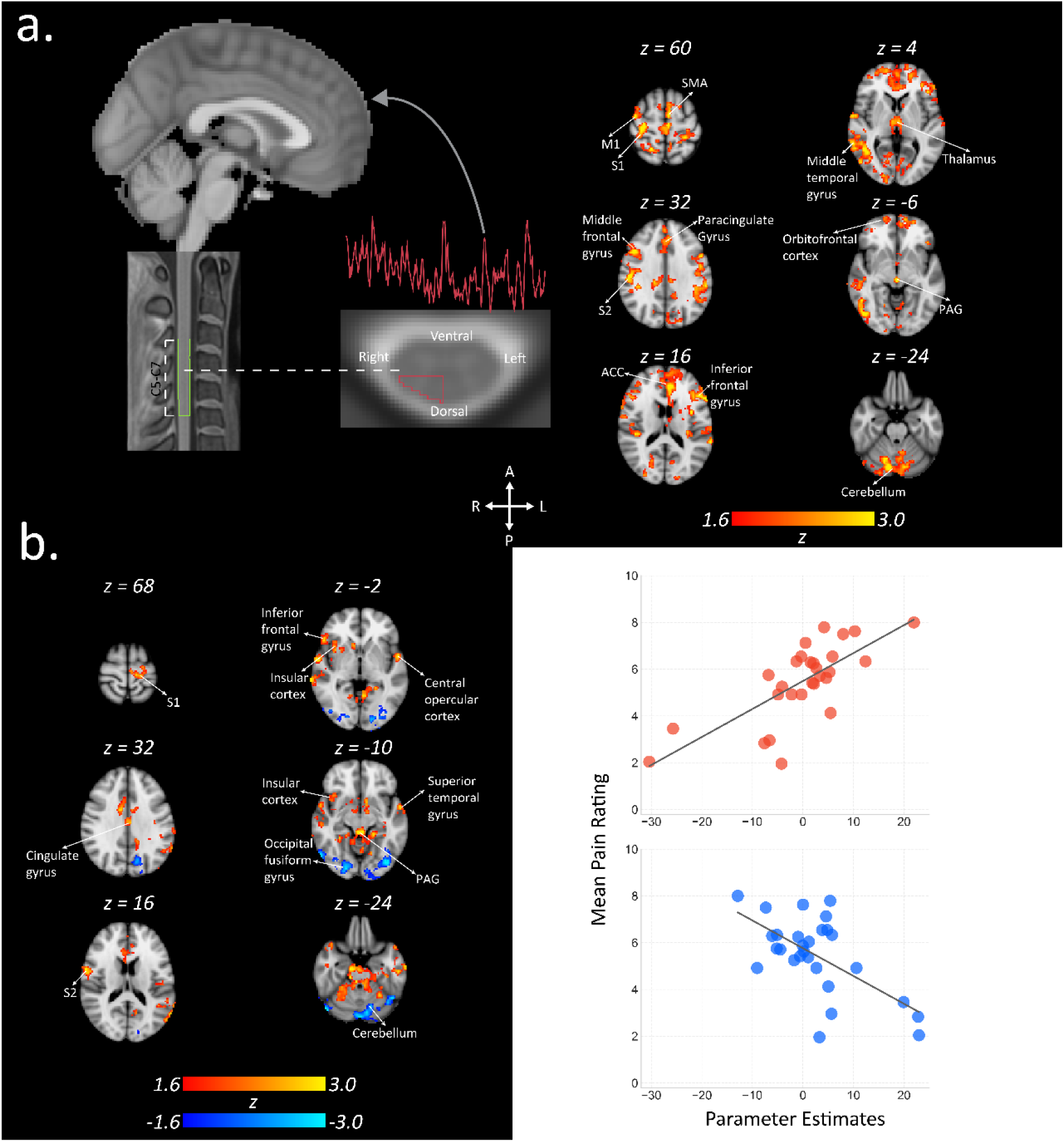
Functional connectivity between the spinal cord and the brain. **a**. Functional connectivity (FC) was calculated between the time-series of the right spinal dorsal quadrant (averaged across segmental levels C5-7) and brain time-series. Coupling between spinal cord time-series and various pain-related brain regions was observed (p < 0.05 FWE-corrected, mixed effects). **b**. Correlation between spinal cord-brain FC strength and mean self-reported pain intensity. Positive correlation between the individual mean pain ratings and the individual strength of FC was observed in distinct pain-related brain regions. (p < 0.05 FWE-corrected, mixed effects). The scatter plots visualize the positive (red) and negative (blue) relationship between parameter estimates of the highlighted regions and mean pain ratings. S1, primary somatosensory cortex; M1, primary motor cortex; SMA, supplementary motor area; S2, secondary somatosensory cortex; ACC, anterior cingulate gyrus; PAG, periaqueductal gray.

Significant positive correlations were observed between spinal cord time-series and pain-related brain regions, including S1, S2, M1, SMA, ACC, orbitofrontal cortex (OFC), middle and inferior frontal gyri, thalamus, paracingulate gyrus, brainstem (PAG), and cerebellum (Fig. 4a, right panel; mixed effects, p < 0.05, cluster correction threshold p < 0.05). No significant negative correlations were detected.

Linking individual variations in spinal–brain FC to behavioral pain ratings revealed a significant positive association in the S1, S2, brainstem (PAG), insula, inferior frontal and superior temporal gyri, and central opercular cortex. Significant negative correlations were observed in the occipital fusiform gyrus and cerebellum (Fig. 4b, left panel; mixed effects, p < 0.05, cluster correction threshold p < 0.05). The scatter plots show that higher parameter estimates in positively correlated regions were associated with higher mean pain ratings, whereas greater activity in negatively correlated regions was associated with lower mean pain ratings (Fig. 4b, right panel).

### Neural activation patterns track temperature in the brain and spinal cord

The activation-based RSA showed that a distributed set of brain regions carried information about temperature (Table 1, Fig. S2 and S3). Importantly, regions whose neural activation patterns tracked subjective pain also overlapped with those tracking temperatures. The activation patterns in PAG and RVM in the brainstem did not track temperature (PAG: t(27) = 2.129, RVM t(27) = −0.523, p > 0.05, n.s) and pain ratings (PAG t(27) = 1.481, RVM t(27) = −1.219, p > 0.05, n.s). Within the spinal cord, activation patterns tracked temperature only in the right dorsal (RD) and right ventral (RV) quadrants (RD: t(27) =4.088, p<0.003; RV: t(27)=3.021, p<0.044, Table 2), consistent with stimulation applied to the right forearm. However, the neural activation in the spinal cord was not predicted by subjective pain ratings (p = xx).

**Table 1.** Brain regions of interest (ROI) where neural activation patterns track temperature and subjective pain representation.

| Brain ROI | Temperature |  |  | Pain |  |  |
| --- | --- | --- | --- | --- | --- | --- |
|  | t | p | Cohen's d | t | p | Cohen's d |
| LH_VisPeri_ExStrInf_1 | 4.648 | * | 0.878 | - | - | - |
| LH_VisCent_Striate_1 | 5.583 | *** | 1.055 | - | - | - |
| LH_VisCent_ExStr_3 | 4.610 | * | 0.8711 | - | - | - |
| RH_VisCent_ExStr_1 | 4.369 | * | 0.826 | 4.650 | * | 0.879 |
| RH_VisPeri_StriCal_1 | 6.804 | *** | 1.286 | 8.115 | *** | 1.534 |
| RH_VisPeri_ExStrSup_1 | 4.453 | * | 0.859 | - | - | - |
| LH_SomMotB_S2_2 | 5.081 | *** | 0.960 | - | - | - |
| LH_SomMotA_1 | 4.884 | *** | 0.923 | 4.277 | * | 0.808 |
| LH_SomMotA_2 | 8.511 | *** | 1.609 | 6.485 | *** | 1.226 |
| RH_SomMotB_Aud_1 | 4.221 | * | 0.798 | - | - | - |
| RH_SomMotB_S2_2 | 6.386 | *** | 1.207 | 4.287 | * | 0.810 |
| RH_SomMotA_3 | 4.364 | * | 0.825 | - | - | - |
| RH_SomMotA_4 | 7.545 | *** | 1.426 | 4.953 | *** | 0.936 |
| LH_DorsAttnA_TempOcc_1 | 4.950 | *** | 0.936 | 4.442 | * | 0.840 |
| LH_DorsAttnB_PostC_2 | 5.413 | *** | 1.023 | - | - | - |
| LH_DorsAttnB_FEF_1 | 7.137 | *** | 1.349 | 5.157 | *** | 0.975 |
| RH_DorsAttnA_TempOcc_1 | 4.489 | * | 0.848 | - | - | - |
| RH_DorsAttnA_SPL_1 | 4.930 | *** | 0.932 | - | - | - |
| RH_DorsAttnB_PostC_2 | 6.085 | *** | 1.150 | - | - | - |
| LH_SalVentAttnA_FrMed_1 | 5.457 | *** | 1.031 | - | - | - |
| LH_SalVentAttnB_PFCmp_1 | 6.113 | *** | 1.155 | 6.679 | *** | 1.262 |
| RH_SalVentAttnA_Ins_1 | 5.105 | *** | 0.965 | 4.828 | *** | 0.912 |
| RH_SalVentAttnA_FrMed_1 | 6.803 | *** | 1.286 | 4.296 | * | 0.812 |
| RH_SalVentAttnB_PFCI_1 | 4.609 | * | 0.871 | 5.185 | *** | 0.980 |
| RH_SalVentAttnB_PFCmp_1 | 4.949 | *** | 0.935 | 5.277 | *** | 0.997 |
| LH_ContA_PFCI_1 | 4.281 | * | 0.809 | 4.263 | * | 0.806 |
| LH_ContA_PFCI_2 | 4.630 | * | 0.875 | - | - | - |
| LH_ContC_pCun_2 | 4.784 | * | 0.895 | - | - | - |
| RH_ContA_IPS_1 | 5.483 | *** | 1.036 | - | - | - |
| RH_ContA_PFCI_1 | 4.450 | * | 0.841 | - | - | - |
| Putamen | 5.656 | *** | 1.069 | - | - | - |

The brain ROI names correspond to Schaefer's parcellation labels. LH: left hemisphere; RH: right hemisphere; ExStrInf: extrastriate inferior; Striate: striate cortex; ExStr: extrastriate cortex; StriCal: striate calcarine; ExStrSup: extrastriate superior; S2: secondary somatosensory; Aud: Auditory; TempOcc: temporal occipital; PostC: post central; FEF: frontal eye fields; SPL: superior parietal lobule; FrMed: frontal medial; PFCmp: medial posterior prefrontal cortex; Ins: insula; PFCI: lateral prefrontal cortex; pCun: precuneus; IPS: intraparietal sulcus. "-" indicated regions are not significant in the corresponding RSA. \* $p < 0.05$ , \*\* $p < 0.01$ , \*\*\* $p < 0.001$ , n.s. $p > 0.05$ , after FWE correction.

**Table 2.** Spinal cord activation patterns track temperature representation.

| Spinal cord regions | Temperature |  |  | Pain |  |  |
| --- | --- | --- | --- | --- | --- | --- |
|  | t | p | Cohen's d | t | p | Cohen's d |
| Right dorsal | 4.088 | ** | 0.773 | 2.461 | n.s | 0.465 |
| Right ventral | 3.021 | * | 0.571 | 1.813 | n.s | 0.343 |
| Left dorsal | 2.817 | n.s | 0.532 | 1.813 | n.s | 0.343 |
| Left ventral | 2.773 | n.s | 0.524 | 1.275 | n.s | 0.241 |

\* $p < 0.05$ , \*\* $p < 0.01$ , n.s. $p > 0.05$ after FWE correction

### Functional connectivity between the brain and spinal cord reflects temperature

The functional connectivity-based RSA showed that temperature significantly predicted connectivity patterns of the right dorsal and right ventral spinal cord with the whole brain (RD: t(27) = 2.510, p < 0.05; RV: t(27) = 3.475, p < 0.002; Fig. 5). By contrast, pain ratings did not explain these connectivity patterns (Table S1). Together, these findings indicate that spinal cord–brain coupling reflects thermal intensity and provide preliminary evidence that the neural representations along the neuroaxis might be primarily organized by sensory temperature cues rather than perceived pain experiences.

**Figure 5.**
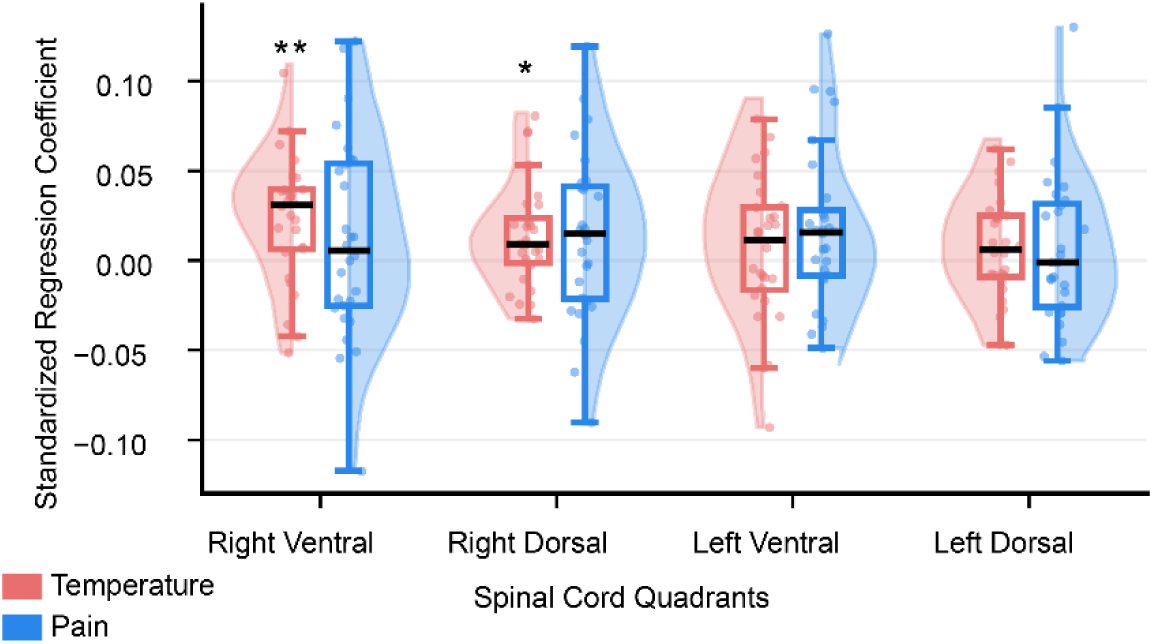
Functional connectivity –based RSA between the cervical spinal cord to the brain. ROI-wise standardized beta regression coefficients are depicted via box plots. For all box plots, the median is denoted by the central mark, and the bottom and top edges of the boxes represent the 25th and 75th percentiles, respectively, with the whiskers encompassing ∼99% of the data. The circles represent individual participants, and half-violin plots show the distribution across participants. The right ventral and the right dorsal quadrants showed functional connectivity RSMs to the brain were predicted by temperature representation (red, RD: t(27) = 2.510, p < 0.05; RV: t(27) = 3.475, p < 0.002), corresponding to the right arm heat stimulation. No spinal cord quadrants with functional connectivity RSMs were predicted by pain rating representations (blue). * p < 0.05, ** p < 0.01.

We subsequently examined whether specific brain networks contributed to this effect using functional connectivity between the right dorsal quadrant and individual brain network. Of the seven parcellated brain networks^39^, no single network exhibited connectivity with the right dorsal spinal cord that was significantly predicted by temperature, while the visual and limbic networks showed trends toward significance (Fig. S4). This could imply that temperature information may be represented in a distributed rather than network-specific manner.

## Discussion

By leveraging simultaneous corticospinal fMRI, this study provides evidence that noxious heat stimulation is encoded not only through local activation across the corticospinal axis but also through their functional coupling. We observed widespread activation in cortical and subcortical regions, as well as in spinal cord segments corresponding to the site of peripheral stimulation. FC analyses further revealed coupling strength between the spinal cord and key pain-processing brain regions positively linked to participants’ individual pain sensitivity. Using RSA, we next showed that multivariate activation patterns in the brain differentially represent temperature and self-reported pain, while the spinal cord specifically encodes information about temperature. Finally, RSA applied to corticospinal FC during heat stimulation showed that these connectivity patterns predominantly represented temperature information in a distributed manner rather than confined to any specific network. This study improves our mechanistic understanding of neural representations of nociception and pain perception along the neuroaxis using corticospinal coupling during noxious heat application.

### BOLD responses to heat

At the level of the brain, pain is largely processed in distributed and bilateral regions^9–11^, including S1, S2, insula, ACC, thalamus, and cerebellum^40,41^. This includes major brainstem regions involved in endogenous descending pain modulation, such as PAG and RVM^42,43^. Using corticospinal fMRI and noxious heat stimulations across three intensities, our study reinforces previous findings^10^ and expands them by showing increased activation in the brainstem and cervical spinal cord. The results indicate a steady increase in the number of both activated and deactivated voxels and corresponding z-scores. Activations clustered in the brain within regions involved in pain processing while deactivations overlapped with the DMN. This pattern confirms previous observations of linear effects across temperatures on brain activity^25,43^. The increase in activated voxels may reflect both recruitment of additional neurons and enhanced spatial spread of cortical/subcortical responses at higher temperatures^27^. This aligns with research that explored brain activation and deactivation upon heat stimulation at two different painful intensities and reported an intensity-dependent (high>low) increase in activation across pain-related regions^44^. In contrast to their original hypothesis, the authors observed larger deactivation across the DMN for lower than higher pain^44^. While others proposed that the DMN consistently displays reductions in activity during a task, independent of its intensity^45,46^, we demonstrate increased deactivation in regions overlapping with the DMN that scales with temperature, as previously reported for certain subregions^11,43,47^. This suggests a shift in neural resources from internally to externally oriented (nociceptive) processing. Interestingly, we also observed an increased number of deactivated voxels in the spinal cord at higher temperatures. While this cannot be interpreted as DMN-related, it may reflect parallel suppression mechanisms or non-specific reductions in neural activity due to high-intensity stimulation.

Early preclinical research by Coghill et al. revealed spinal responses to innocuous and noxious heat stimulation of rat hind limbs, with greater nociceptive processing detected at higher temperatures^48^. In correspondence with neuroanatomy, the activity was mainly observed in the ipsilateral dorsal horn but to some degree also in the contralateral dorsal horn and ventral horns, especially at higher temperatures. Further, using in vivo two-photon calcium imaging in mice, Ran et al.^49^ captured graded heat-evoked responses in the spinal cord and found neurons in the dorsal horn to represent a range of heat stimulation intensities. In agreement, we detected increased spinal activation and a broader spatial distribution at higher temperatures, suggesting graded nociceptive transmission up the spinothalamic and spinoreticular tracts. The activation patterns extended into intermediate and ventral regions which may reflect the co-activation of spinal interneurons and motoneurons involved in executing protective reflexes such as the flexion response^13,17^, though further studies with higher sensitivity are needed to confirm this interpretation.

Further, the activation cluster located in the medioventral spinal cord can most likely be attributed to the sulcal vein draining the ventral GM horns^50^.

### Functional coupling along the central nervous system

To explore the intricate interplay between spinal and supraspinal networks during nociceptive processing, we correlated spinal cord and whole-brain time-series, extending the approach of Sprenger et al who limited their analysis to predefined brain ROIs^24^. They highlight positive FC between the spinal cord and S1, S2, insula, striatum, midbrain, amygdala, and hypothalamus, and spinal cord-PAG coupling strength was positively correlated to the mean pain rating^24^. We observed coupling between the ipsilateral spinal dorsal quadrant and additional key brain regions involved in pain processing, including motor regions, ACC, paracingulate gyrus, OFC, frontal and temporal gyri, thalamus, and cerebellum. Previous research showed coupling between bilateral posterior cerebellum and dorsal horns of the cervical spinal cord during rest^51^. Positive relationships between mean self-reported pain intensity and individual strength of FC between the spinal cord and brain expanded beyond the PAG^24^ to S1, S2, central opercular cortex, insula, and frontal and temporal gyri. This expands previous studies showing higher pain perception to be reflected by stronger brainstem-cortical coupling^52^. Parts of the occipital fusiform gyrus and cerebellum comprised negative relationships between FC strength and self-reported pain, possibly reflecting anti-correlated sensory suppression consistent with the role of these areas in sensory filtering and emotional regulation^53^. Altogether, this highlights the multifaceted functional relevance of the spinal cord-brain interplay during nociceptive processing.

While classically being thought to be exclusively involved in motor processing and coordination, the cerebellum was shown to receive noxious inputs from the spinal cord and be capable of modulating nociceptive processing via higher-order areas^54–57^. In line, previous fMRI studies observing differential cerebellar activation patterns for noxious vs. innocuous stimulations may reflect not only somatosensory processing but also pain intensity perception^58,59^. Our results show bilateral cerebellar activation at the lowest noxious temperature, which spans almost all of the cerebellum at higher temperatures. Further, positive findings of spinal cord-cerebellum FC and its correlation with self-reported pain reinforce its role in nociceptive processing via the spinocerebellar pathway and suggest an involvement in the subjective experience of pain, either directly or via cognitive cortical/sub-cortical areas. During rest, pain intensity was shown to be negatively correlated with FC strength between the cerebellum and the DMN in adolescents with pain^60^, highlighting the cerebellum’s crucial role in both acute and chronic pain processing.

Our findings, together with previous corticospinal fMRI research^24^, showcase the critical role of spinal-brainstem interactions relaying aversive physical stimuli to be integrated into the multifaceted pain perception process at the brain level^61,62^. Strong preclinical^63–67^ and clinical^15,20,62,68–70^ evidence supports the role of the PAG and RVM as key brainstem nuclei involved in pain processing and endogenous pain modulation; its magnitude being dependent on the intensity of stimulation. Using combined brainstem–spinal cord fMRI, studies have highlighted varying FC changes related to differences in pain sensitivity^71^ and enhanced FC for noxious vs innocuous heat stimulation^72^. This confirms our observations of a positive coupling between the right spinal dorsal horn and the PAG and its strength being correlated with subjective pain ratings. Further, spinal connectivity with prefrontal regions (OFC, inferior frontal gyrus) indicates engagement of the descending pain-modulatory system, and association with pain ratings suggests inter-individual variability in top-down control efficacy^14^.

### Brain and spinal cord activation and coupling differentially track temperature and self-reported pain

It has been well established by multivariate analytical approaches that nociceptive input and subjective pain experience are encoded in distributed patterns of brain activity rather than solely in changes of signal amplitude^26–28^. However, similar frameworks have yet to be extended to the spinal cord, leaving open how integrated corticospinal activity patterns jointly encode stimulus intensity and perceived pain. To the best of our knowledge, we are the first to apply RSA^29^ to simultaneous spinal cord-brain fMRI to explore how corticospinal coupling reflects representations of nociception and self-reported pain, leveraging the multivariate approach to better explain the large intra-^13^ and inter-individual^18^ variability underlying fMRI signal. Consistent with prior brain literature^26–28^, our RSA results based on regional activations showed widespread activation patterns organized by temperature.

Interestingly, a subset of these brain regions also encoded subjective pain, mostly those within salience/ventral attention and sensorimotor networks, including insula and PFC^73^. This aligns with the literature highlighting the important roles of these regions in processing salient stimuli and endogenous pain modulation^62,74^.

The right dorsal and right ventral quadrants of the spinal cord tracked temperature but not pain. Our results also highlighted that corticospinal FC during heat stimulation predominantly represented temperature information in a distributed manner. Notably, the PAG also demonstrated significant temperature encoding before correction for multiple comparisons, supporting our spinal-PAG coupling results in the FC analysis, and the critical role of the brainstem as part of a spino-bulbo-spinal pain feedback loop^64^. Together, our findings underscore the importance of measuring spinal cord activity to better understand the pain-modulating pathway along the neuroaxis. They provide preliminary evidence on how both distinct and overlapping neural mechanisms underlie the nociceptive and pain processing^28^. Future studies with a more fine-grained heat stimulation design could further delineate the gradient representations of these dimensions of pain along the corticospinal neuroaxis.

Several limitations should be discussed. First, as in other spinal cord fMRI studies employing GE-EPI acquisitions^75–79^, BOLD responses in the spinal cord are not confined to gray matter but extend toward the cord edge due to signals from draining veins^80,81^. While excluding the signals from outside the cord could help reduce these effects, spatial smoothing may still blur peripheral vascular responses into the cord, compromising spatial specificity. Second, spinal cord findings were thresholded more liberally compared to brain, given low SNR, small spatial extent, and substantial physiological noise, a thresholding approach not uncommon in spinal cord fMRI literature^13,78,82^. While the use of mixed-effects models and known anatomical priors partially mitigates this issue, results should be considered preliminary and replicated with larger samples and optimized acquisition pipelines. Finally, task contrasts and FC estimates are correlational and cannot establish directionality or causality. Cognitive modulators of pain (e.g., attention, expectation, affect) were not measured, limiting interpretation of FC and potential descending effects. Future work incorporating explicit cognitive modulation paradigms is needed.

In conclusion, this study provides compelling evidence that noxious heat engages nociceptive pathways across the CNS in an intensity-dependent manner, reflected in local activation and functional coupling across brain, brainstem, and spinal cord. Using simultaneous corticospinal fMRI, we show that increasing temperature leads to graded recruitment of bilateral brain regions and engagement of descending modulatory circuits. Neural activation predominantly within sensorimotor and salience networks also scales with subjective pain, revealing distinct processes underlying nociception and pain perception.

These results highlight the translational potential of simultaneous brain–spinal cord imaging for identifying systems-level biomarkers of stimulus intensity and self-reported pain encoding. They provide a foundation to be explored in patient populations when testing pathological spinal cord-brain connectivity in chronic pain and mediating effects of pharmacological and non-pharmacological interventions.

## Data and code availability

The data can be made available upon request; however, due to ongoing analyses, they are not publicly accessible at this time. The code to process and analyze the images can be accessed at https://github.com/yiyuwang/ThermalEncoding_BrainSpinalCord

## Author contributions

D.P.: Methodology, Formal Analysis, Visualization, and Writing (Original draft, Review & Editing).

Y.W.: Methodology, Formal Analysis, Visualization, and Writing (Original draft, Review & Editing).

M.K.: Methodology, Formal Analysis, Visualization, and Writing (Original draft, Review & Editing).

C.L.: Conceptualization, Data Acquisition, Methodology, Pre-processing, Funding Acquisition, Supervision, and Writing (Review & Editing).

J.F.: Writing (Review & Editing)

T.D.: Writing (Review & Editing)

V.O: Writing (Review & Editing)

T.I: Writing (Review & Editing)

S.S.: Methodology and Writing (Review & Editing)

T.S.: Methodology and Writing (Review & Editing)

P.L.: Methodology and Writing (Review & Editing)

M.T.: Methodology and Writing (Review & Editing)

K.W.: Conceptualization, Methodology, Formal Analysis, Funding Acquisition, Supervision, and Writing (Review & Editing).

G.G.: Conceptualization, Methodology, Funding Acquisition, Supervision, and Writing (Review & Editing).

S.M: Conceptualization, Methodology, Funding Acquisition, Supervision, and Writing (Review & Editing).

## Declaration of competing interests

The authors do not have any competing interests to declare.

## Acknowledgements

We thank all the volunteers for participating in this study.

## Funding

This study was supported by grants from the National Institute of Neurological Disorders and Stroke (grant numbers R01 NS109450, K23NS104211, L30NS108301, R01NS128478, and K24NS126781) and the National Institute on Drug Abuse (T32DA035165). The content is solely the responsibility of the authors and does not necessarily represent the official views of the National Institutes of Health. Dario Pfyffer is supported by a Swiss National Science Foundation Postdoc.Mobility Fellowship grant (P500PM_214211). Valeria Oliva is supported by the Italian National Institute of Health (Starting Grant for Young Researchers CUP I85E23001090005).

## Notes

### Competing Interest Statement

The authors have declared no competing interest.

