## Supplemental material for "Differential encoding of noxious heat and self-reported pain along corticospinal networks: a simultaneous spinal cord-brain fMRI study"

### Supplementary material:

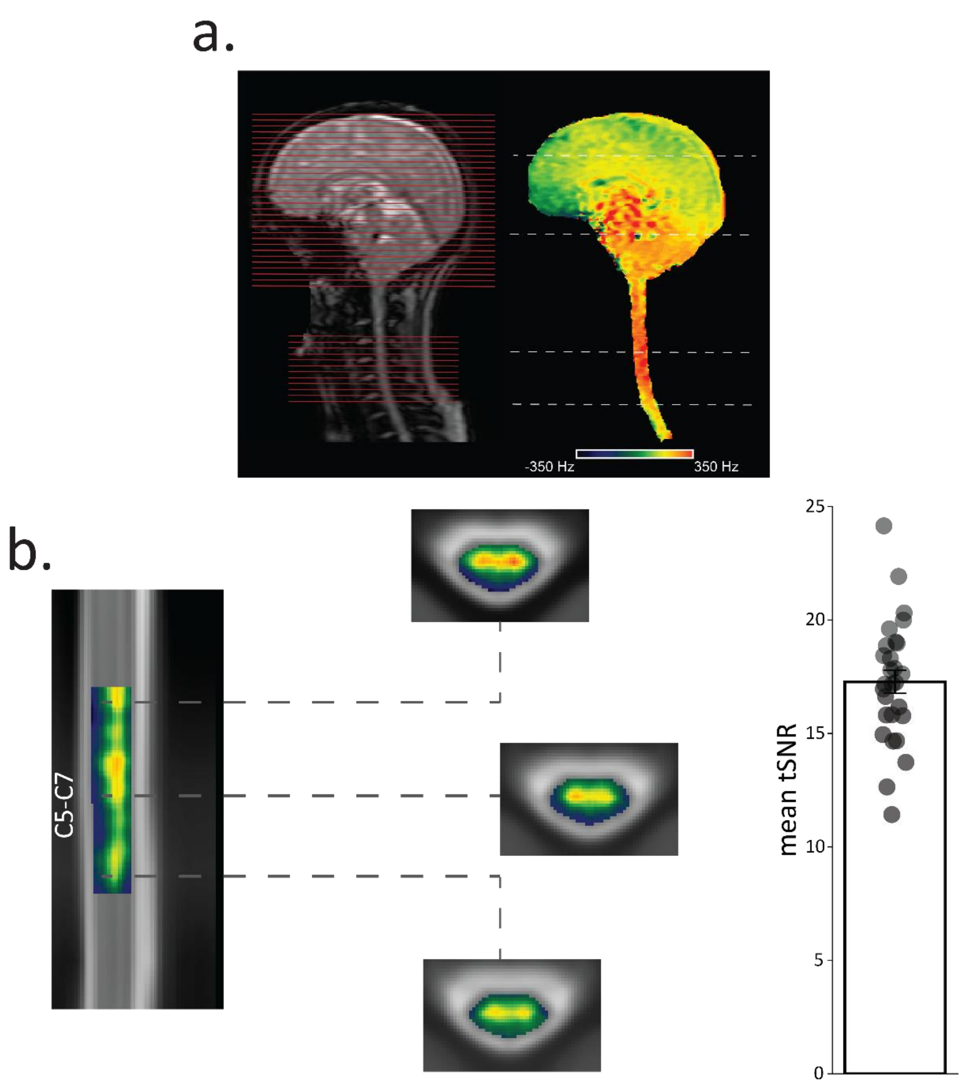


***Figure S1.*** **Corticospinal functional magnetic resonance imaging (fMRI) data acquisition and temporal signal-to-noise ratio (tSNR). a**. An exemplary corticospinal imaging acquisition for a participant is depicted. A sagittal localizer image shows the position of slices (red lines) for the brain and spinal cord. On the left panel, a sagittal field map is masked to depict field variations in the tissues of interest. Slice-wise shimming was employed as explained previously. **b**. Spinal tSNR map. Group-average coronal tSNR map after preprocessing and three exemplary sagittal slices are shown in PAM50 template space (tSNR range from 10 to 20). The bar graph shows the tSNR of spinal cord for individual subjects.

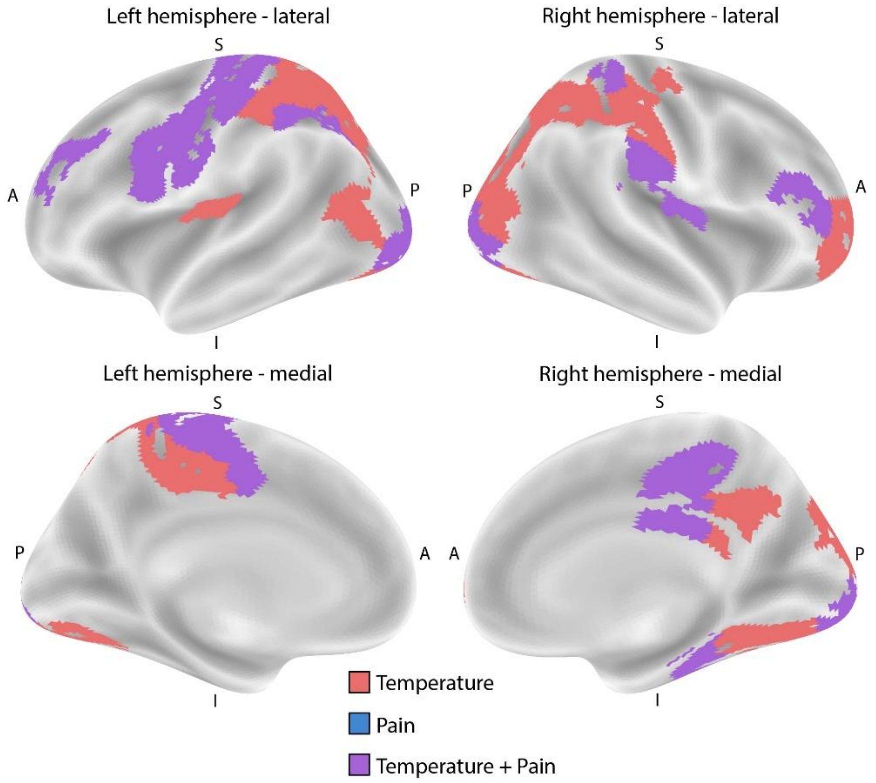


***Figure S2. Surface plots of brain regions for temperature and pain encoding.*** Brain regions with activation patterns predicted by temperature RSMs (red) or pain (blue, no results survived multiple comparison). Brain regions with activation patterns predicted by both the temperature and pain rating RSMs are shown in purple.

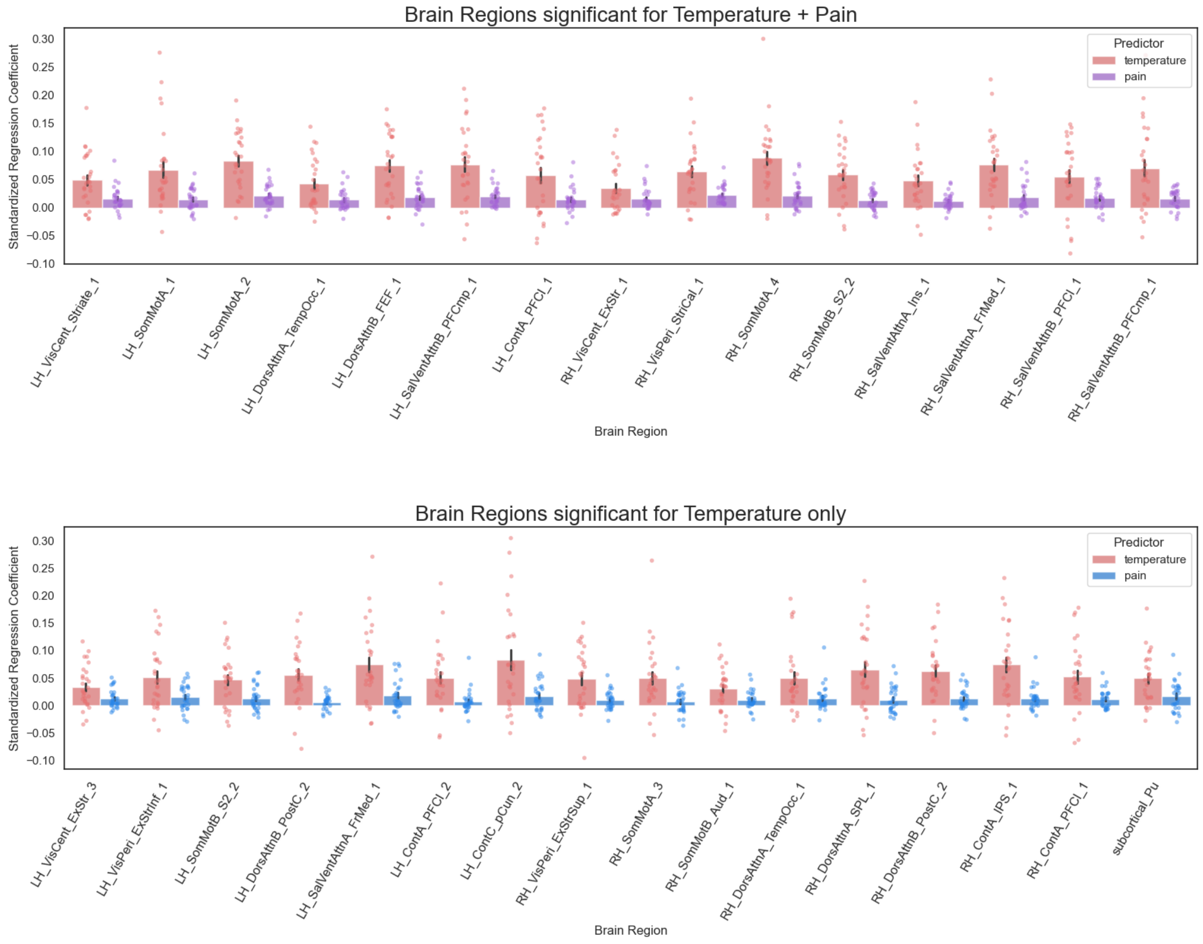


***Figure S3. Standardized regression coefficients of brain regions for temperature and pain encoding.*** Brain regions tracking both temperature and pain ratings (upper panel) and brain regions tracking temperature only (lower panel). The neural activity RSMs of these regions were not predicted by pain ratings.

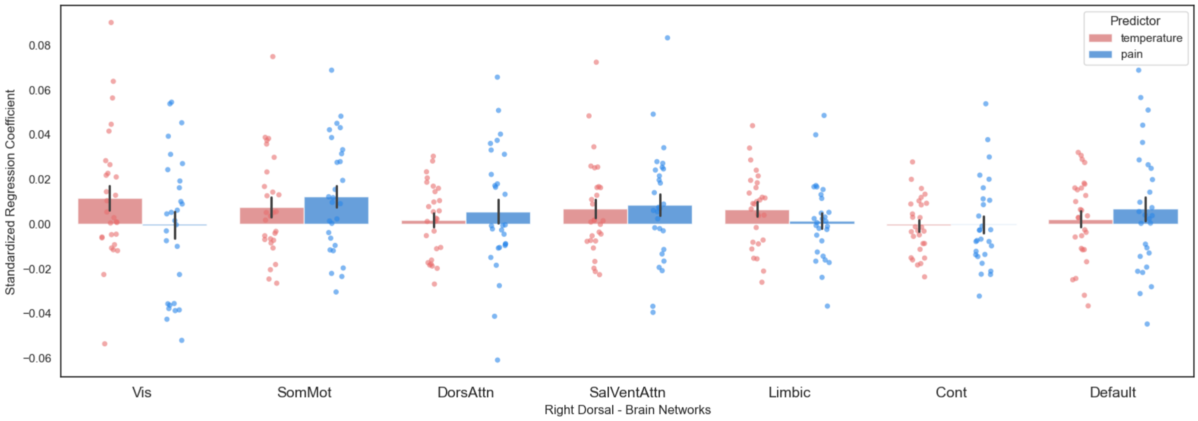


***Figure S4. Spinal cord-brain network coupling for temperature and pain encoding.*** The functional connectivity between the right dorsal quadrant to individual brain networks was not predicted by temperature or pain, suggesting the results were not likely to be driven by a single neural network. (Vis: Visual; SomMot: Somatomotor; DorsAttn: Dorsal Attention; SalVentAttn: Salience Ventral Attention; Cont: Control; Default: Default Mode).

Table S1. Spinal cord-brain connectivity patterns track temperature representation.

| Spinal cord regions | Temperature | | | Pain | | |
|---|---|---|---|---|---|---|
| | t(27) | p | Cohen's d | t(27) | p | Cohen's d |
| Right dorsal | 2.510 | * | 0.474 | 1.802 | n.s | 0.341 |
| Right ventral | 3.475 | ** | 0.657 | 1.152 | n.s | 0.218 |
| Left dorsal | 1.218 | n.s | 0.230 | 0.755 | n.s | 0.143 |
| Left ventral | 0.770 | n.s | 0.146 | 2.027 | n.s | 0.230 |
